# Buffaloed in Brandenburg: Germany’s first Brush with Foot-and-Mouth Disease after four Decades of Freedom

**DOI:** 10.64898/2026.03.30.713672

**Authors:** Michael Eschbaumer, Christoph Staubach, Florian Pfaff, Jörn Gethmann, Katja Schulz, Lisa Rogoll, Sabine Bock, Wulf-Iwo Bock, Christoph Schulze, Ronny Marquart, Nicole Reinhardt, Stephan Nickisch, Nadine Kakerow, Sabrina Freter, Annett Rudovsky, Kerstin Albrecht, Sandra Leo, Christina Haarmann, Sarah Lenz, Barbara Hoffmann, Sten Calvelage, Dirk Höper, Patrick Zitzow, Angele Breithaupt, Can Çokçalişkan, Ünal Parlak, Sharon Karniely, Laith Mohammed Salih Abdulrasool, Nick Knowles, Guillaume Girault, Aurore Romey, Labib Bakkali Kassimi, Donald P. King, Christa Kühn, Carola Sauter-Louis, Martin Beer

## Abstract

Foot-and-mouth disease (FMD) virus is one of the most feared and most consequential pathogens of livestock worldwide. It can be spread rapidly by the transboundary movement of animals, animal products and byproducts. In January 2025, Germany detected its first FMD outbreak since 1988 in extensively reared water buffalo on a small farm in the state of Brandenburg, directly outside Berlin, the federal capital. Immediate control measures including a standstill for movements of susceptible animals and pre-emptive culling were implemented by the veterinary authorities. Whole-genome sequencing identified the virus as serotype O, topotype ME-SA, lineage SA-2018 and revealed extensive recombination, but cross-neutralization assays suggested good heterologous protection by an O/PanAsia-2 vaccine strain. Epidemiological back-calculation placed the time of virus introduction in late December 2024. Although the entry route remains unresolved, human-associated introduction is most likely. Network analysis revealed minimal farm connectivity, and simulations predicted low potential for onward transmission, which is consistent with the outbreak being ultimately restricted to a single herd. This event underscores the constant and unpredictable risk of introduction of the virus. Early detection through increased awareness and comprehensive differential diagnostics as well as the international collaboration of veterinary services, laboratories and experts are essential in the face of the global presence of FMD.

## INTRODUCTION

Foot-and-mouth disease (FMD) is a highly contagious viral disease of cloven-hoofed animals, characterized by fever, vesicular lesions on the mouth, feet and udder, and severe economic consequences for agriculture worldwide^1^. The causative agent, FMD virus (FMDV; *Aphthovirus vesiculae*, family *Picornaviridae*), is endemic in Africa and large parts of Asia, but was last reported in Germany in 1988^2^. FMDV has a single-stranded, positive-sense RNA genome which encodes a polyprotein that is cleaved into structural and non-structural sub-proteins which are essential for its life cycle^3^. The capsid protein VP1 is particularly important as it carries the major antigenic sites that trigger immune recognition and play a vital role in attaching to host cells^4^.

Any outbreak of FMD not only carries the risk of rapidly spreading infection and disease in ungulates, but also precipitates drastic control measures and dramatic economic losses^5^. For example, the major FMD epizootic in the United Kingdom in 2001 culminated in the culling of millions of animals and caused billions of dollars in damage across several sectors of the economy^6^. FMD outbreaks in free regions are always serious, but Germany’s first case in nearly forty years turned out to be exceptional in several respects.

## RESULTS AND DISCUSSION

### The outbreak

In early January 2025, after three of 14 water buffalo (*Bubalus bubalis*) in a grazing herd in Brandenburg, Germany, had died within two days, the owner took the carcass of the third animal (ear tag 334) to the state veterinary diagnostic laboratory for a post-mortem examination. Upon observing expansive erosions of the oral and ruminal mucosa (including approximately one-third of the lingual epithelium), pulmonary oedema and necrotizing myocarditis, the state laboratory performed standard differential diagnostics^7^ using tissue samples taken at necropsy. Lung tissue taken at necropsy was highly positive for FMDV RNA by real-time RT-PCR (Cq 25). The samples were immediately sent to the German national reference laboratory (NRL) for FMD, which corroborated the earlier PCR results and confirmed the outbreak on 10 January (see timeline in **Figure 1**). After immediate notification in the European Union Animal Disease Information System (ADIS) and the World Animal Health Information System (WAHIS), the World Organisation for Animal Health (WOAH) suspended Germany’s FMD-free status, effective from 9 January 2025.

**Figure 1:**
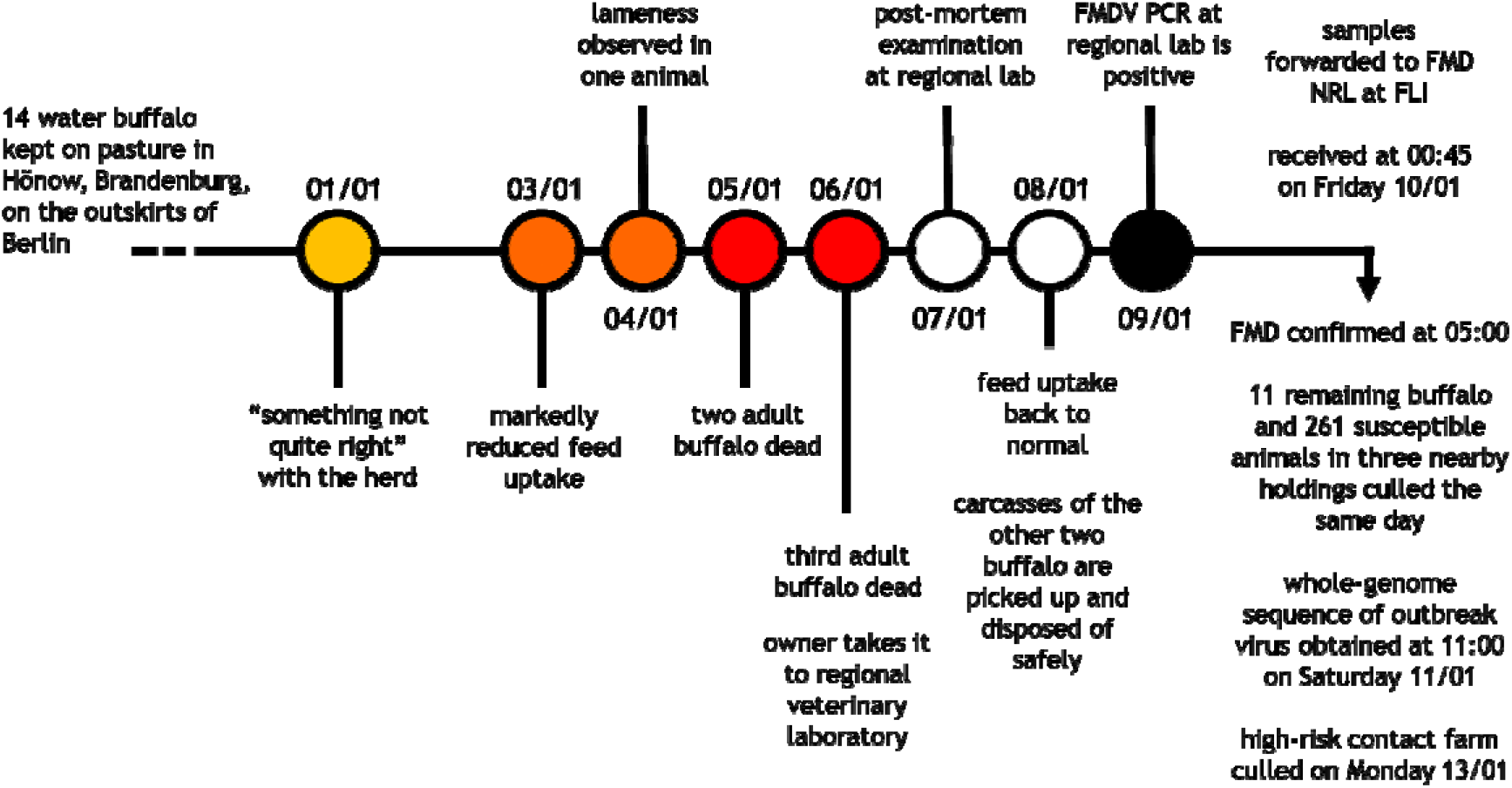
Timeline of events, clinical signs and FMD diagnostics in Germany in January 2025.

The veterinary authorities in Brandenburg immediately took action and the remaining 11 buffalo were culled the same day, followed by 261 susceptible animals (245 pigs, 13 sheep and 3 goats) in three nearby holdings and 61 susceptible animals (3 cattle, 39 sheep, 19 goats) at a high-risk contact farm that had received hay from the infected farm. All culled ruminants and 189 of the 245 culled pigs were tested for FMDV.

The carcasses of the first two buffalo in the index herd, which had died on 5 January, had already been disposed of without sampling. No FMDV-specific antibodies were detected in the third dead buffalo (ear tag 334), but the remaining 11 animals in the index herd were all positive for antibodies to both non-structural and structural proteins of FMDV and FMDV RNA at the time of culling. Clinical examinations of the 11 culled buffalo revealed oral (in 9 animals) and/or podal (in 2 animals) lesions typical of FMD, which were already healing (**Figure 2**). One animal (ear tag 165) had no conspicuous lesions in the mouth or on the feet. The oldest lesions (found in animal 565) were determined to be clearly older than one week at the time of culling on 10 January. High levels of FMDV RNA (Cq 21–23) were detected in the podal lesions, but the oral lesions and nasal swabs were only weakly positive (Cq >31). Only 4 animals in the herd had detectable FMDV RNA in serum (see **Supplemental Table S1**). Intralesional FMDV antigen and virus genome was detected in the heart of buffalo 334 by immunohistochemistry and RNA *in situ* hybridization, respectively (see **Supplemental Figure S1**). Thus, the necrotizing myocarditis was most likely induced by FMDV infection and interpreted as the cause of pulmonary oedema and death.

**Figure 2:**
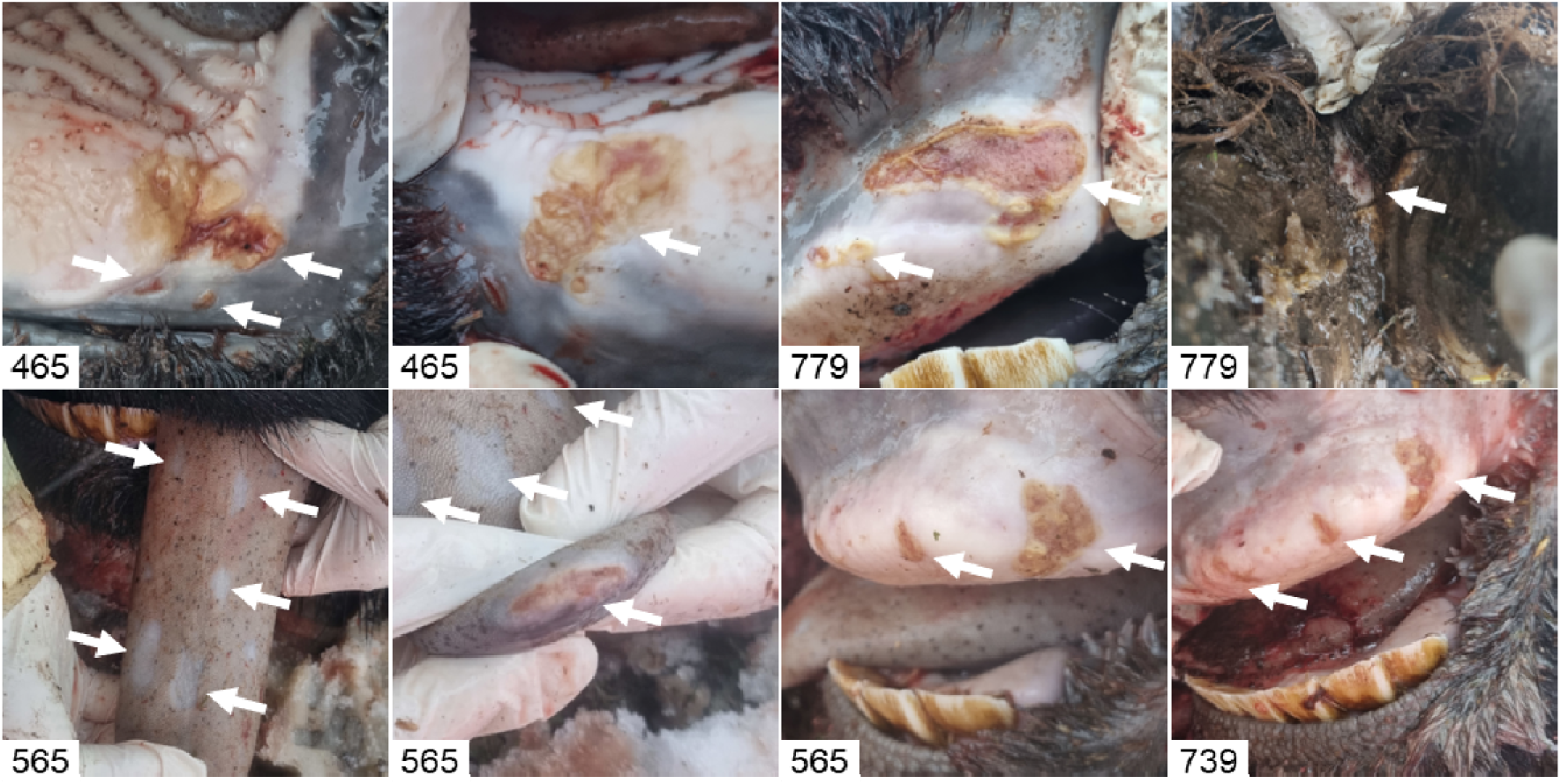
Lesions on the gum, tongue and in the interdigital cleft of culled water buffalo on the affected farm. Pictures are labelled with the last three digits of the animal’s ear tag number. Lesions are marked with white arrows. The oldest lesions were found on the tongue of animal 565 (bottom left).

No FMDV RNA or specific antibodies were found in any animal of the other four depopulated holdings.

### Genetic origin of the German FMDV

Based on the coding region of the viral capsid protein VP1 (633 nucleotides), the virus isolated from buffalo 334 was identified as FMDV serotype O, topotype ME-SA and lineage SA-2018 (**Supplemental Figure S2a**). The most closely related VP1-coding sequence that was available for comparison corresponded to a virus isolated in late November 2024 in far eastern Türkiye along the border with Armenia, Azerbaijan and Iran. These viruses were sampled 43 days apart from each other and shared 632 of 633 nucleotides (nt) of their VP1 sequence. Since in lineage SA-2018 the genetic drift of VP1 is strongly correlated with the sampling date (**Supplemental Figure S2b**), we conducted a time-aware phylogeny analysis to gain insights into the genetic origin of the FMD virus responsible for the outbreak in Germany (**Figure 3a**). The SA-2018 lineage was first reported in India in 2018^8^, but was later found in Sri Lanka, Nepal, Bangladesh, Oman, and the United Arab Emirates^9^. The VP1 sequence of the virus detected in Germany belongs to a genetically distinct sub-lineage that was first recorded in 2022 in Bangladesh, India and Nepal. Between 2023 and 2025, different genetic clades of this sub-lineage were also detected in Israel, Iran, Iraq, Syria and Türkiye (**Figure 3b, c**). Interestingly, the phylogenetic and temporal structure of the relationships between these VP1 sequences showed that genetically distinct variants were present at roughly the same time in 2025, e.g., in Türkiye and Israel, supporting multiple introduction events into Western Asia. Later in 2025, genetically similar variants of the virus detected in Germany were also found in Israel and in additional FMD cases in Türkiye, indicating at least temporary infection chains in these geographic regions.

**Figure 3:**
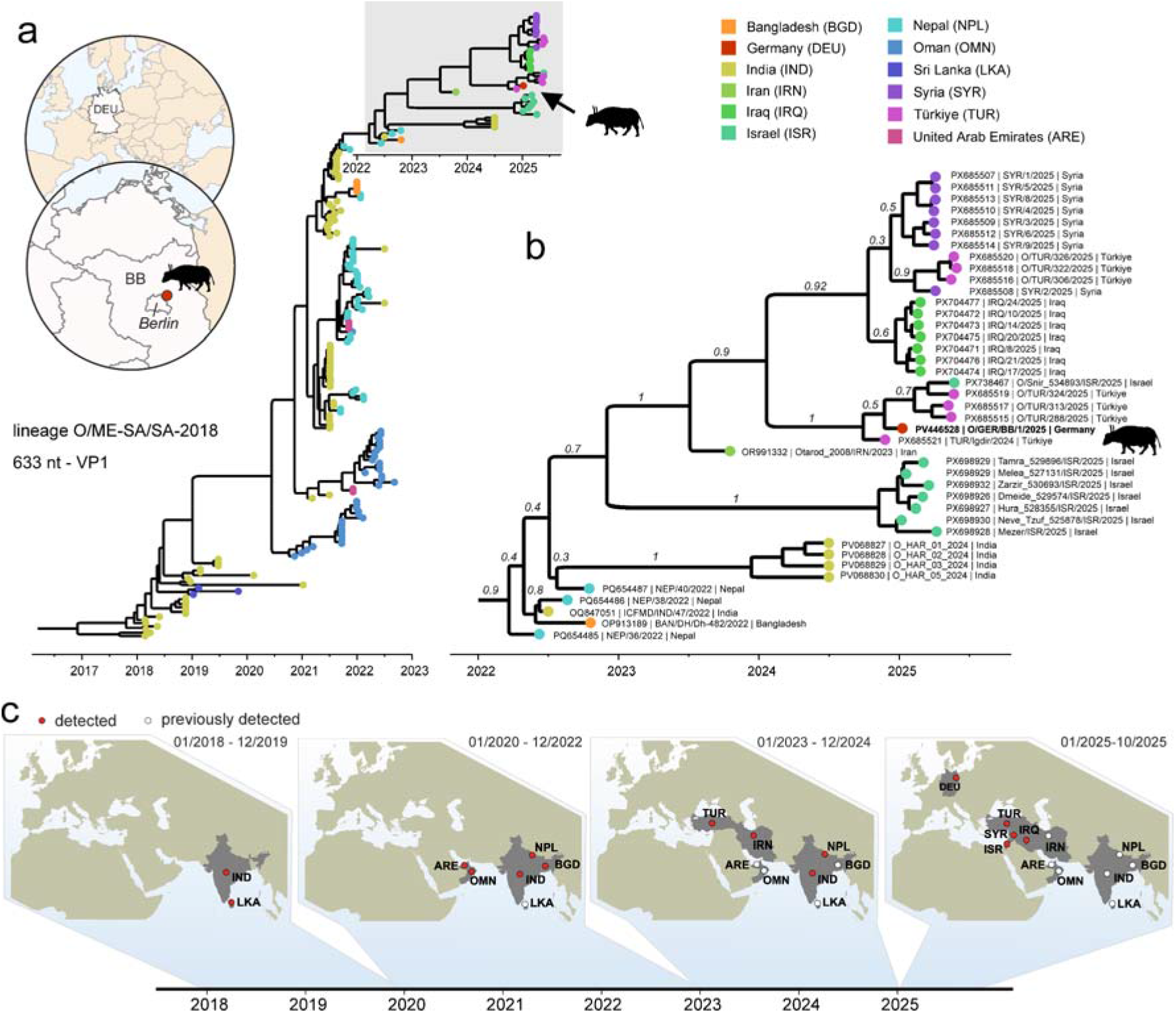
Genetic origin of the German FMDV strain. **a** FMDV was detected in January 2025 in a herd of water buffalo in the German state of Brandenburg, close to Berlin. Time-aware phylogenetic analysis of the VP1-coding region revealed the virus to be of lineage O/ME-SA/SA-2018. **b** The closest genetic and temporal relatives were found in Türkiye and Israel. **c** Timeline showing the geographic spread of the SA-2018 lineage. The SA-2018 lineage was initially identified in India in 2018, and until 2022 was detected in Sri Lanka, Nepal, Bangladesh, Oman and the United Arab Emirates. By the end of 2024, the lineage had been spread to Iran and Türkiye, and from 2025 onwards outbreaks from Iraq, Israel, Syria and Germany were reported.

The number of full FMDV genome sequences from lineage SA-2018 available for our analysis is currently low (n = 12) and limited in both time and geographic origin, and full genome comparisons are further complicated by frequent genome recombination events^10^. However, the comparison of the German FMDV genome to its closest genetic relatives showed that the virus has undergone at least one or likely multiple recombination events with viruses from lineages O/ME-SA/Ind-2001, O/ME-SA/PanAsia-2, A/ASIA and Asia1/ASIA (**Supplemental Figure S3**). While FMDV genomes from lineage SA-2018, sampled between 2019 and 2022 in Bangladesh, Nepal, Sri Lanka, and the United Arab Emirates, were genetically highly similar, only parts of the German FMDV genome were phylogenetically closely related to these viruses (**Figure 4a**). In detail, the region spanning the 5’ UTR to 2A, including the VP1 coding region, phylogenetically groups with the other viruses from lineage SA-2018. The genomic region spanning 2B to 3B of the virus detected in Germany, however, phylogenetically grouped with viruses from the O/ME-SA/Ind-2001 lineage. Regions 3C, 3D and the 3’ UTR do not match the SA-2018 lineage viruses and indicate recombination with previously detected viruses from lineages O/ME-SA/PanAsia-2, A/ASIA and Asia1/ASIA (**Figure 4b**).

**Figure 4:**
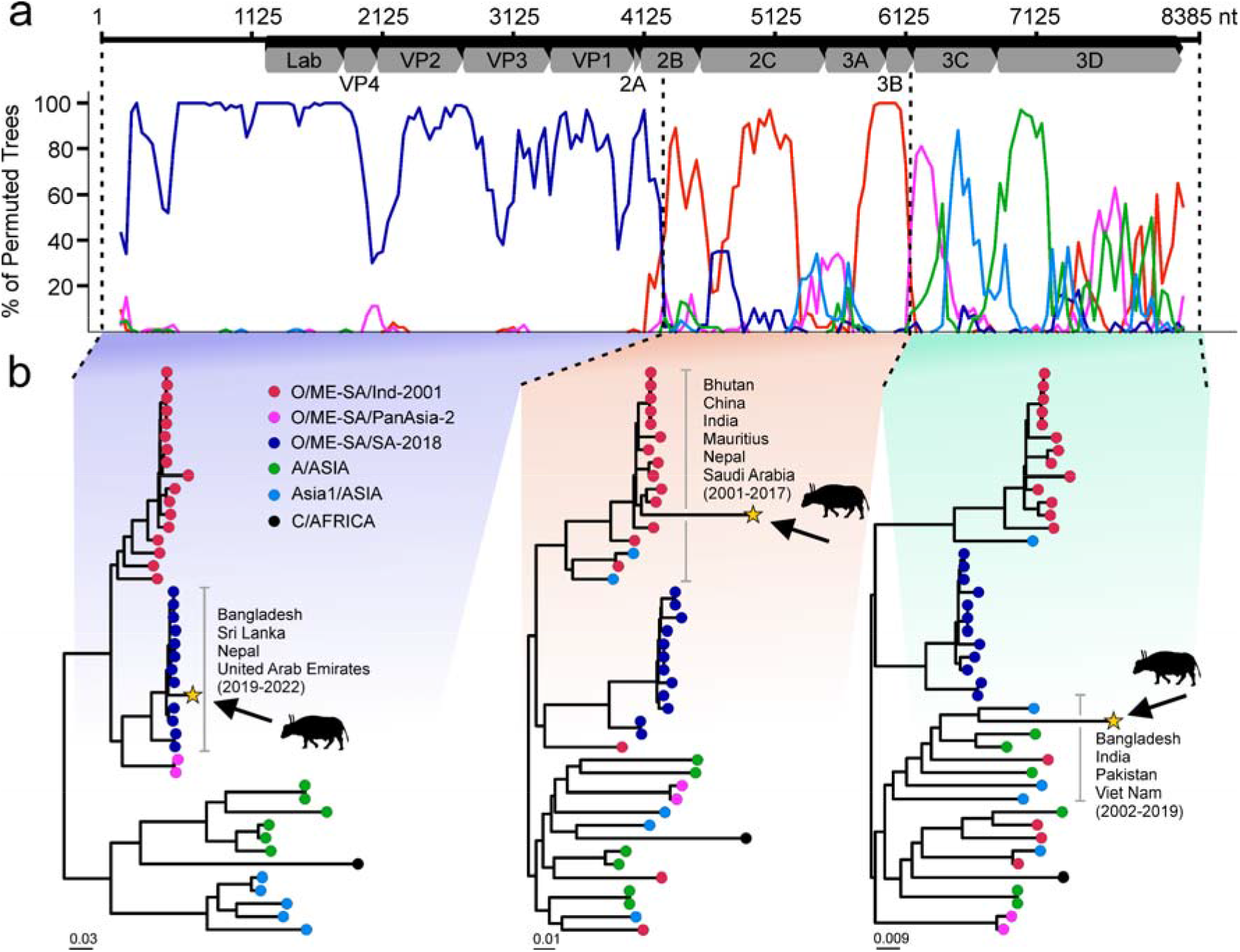
Signals of recombination in the German FMDV strain. **a** A sliding-window approach (BootScan) was used to detect recombination events in an alignment of selected full FMDV genome sequences. The phylogenetic position of the German FMDV in relation to the references was observed in each window and cumulated for 100 bootstraps. Reference sequences were selected by querying 500-nt windows of the FMDV O/GER/BB/1/2025 genome against public databases and aligning the top full-genome hits together with all available SA-2018 lineage genomes. Each coloured line shows the percentage of permuted phylogenetic trees in which O/GER/BB/1/2025 clusters with a given sequence group (see colour legend in panel b) across a sliding window. **b** Maximum-likelihood trees of three genome regions highlight the complex genetic makeup of the FMDV strain detected in Germany.

### Vaccine matching

An isolate of the German FMD virus was used for *in vitro* vaccine matching using homologous and heterologous virus neutralization tests. Three vaccine strains available from Boehringer Ingelheim (O-3039, O PanAsia-2 and O_1_ Manisa) and the corresponding bovine vaccinal sera (BVS) were considered. On average, the highest r_1_-values and heterologous titres, which are considered the best predictors of cross-protection *in vivo*, were obtained with the O PanAsia-2 and O_1_ Manisa vaccines (see **Supplemental Table S2**). The FMD vaccine bank of the German federal states was activated and 750,000 doses of O PanAsia-2 vaccine were produced. Ultimately, the decision was made not to vaccinate, as the epidemiological situation did not warrant it.

### Epidemiology, network analysis and modelling

The outbreak farm was located east of Berlin, in a region of low farm and livestock density (see **Figure 5** and **Supplemental Figure S4**). It was a breeding farm, which consisted of one bull, six cows, and seven calves, kept on a partially fenced pasture of 8.5 hectares. Minor behavioural changes and a reluctance to feed were first observed on 1 January 2025. Feed intake was markedly reduced on 3 January and one calf was lame the following day (see timeline in **Figure 1**). The ear tag number of this animal was not recorded.

**Figure 5:**
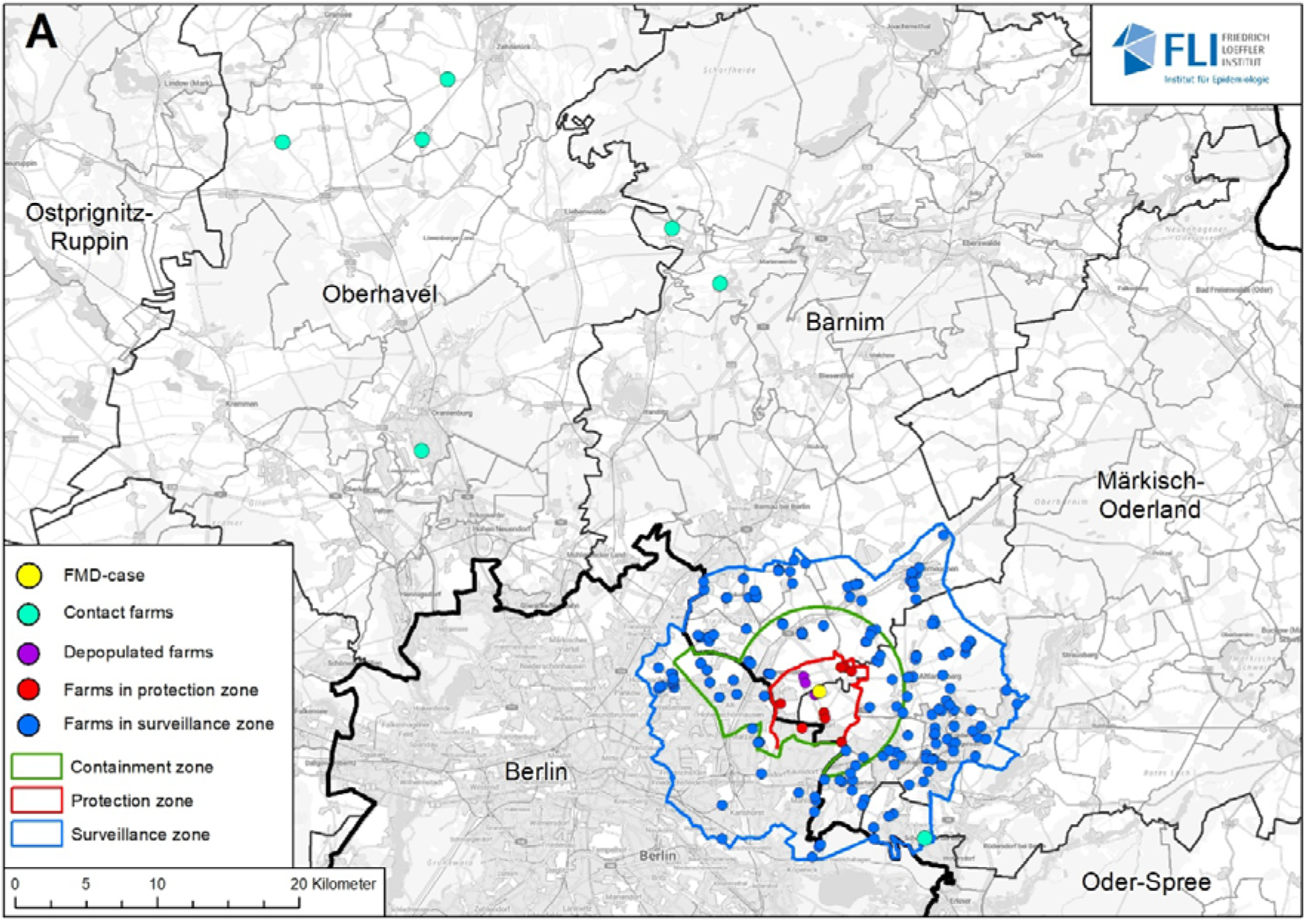
Location of farms in restricted zones as well as contact farms. The red and blue lines depict the protection and surveillance zone around the outbreak farm. The green line shows the containment zone, which was approved by WOAH on 12 March 2025. The 7 farms shown in turquoise are contact farms. The contact farm south of the outbreak farm was classified as high risk, as it had received hay from the outbreak farm, while the other contact farms (north-west of the outbreak farm) were linked through the rendering vehicle.

The carcasses of the first two dead buffalo were collected by a rendering vehicle on 8 January without any samples being taken. The rendering vehicle subsequently stopped at six more farms on the same day, which were considered contact holdings in the subsequent epidemiological investigation.

Assuming an incubation period of approximately 3–7 days and the development of antibodies around 7–10 days after infection, it can be concluded that the virus was introduced to the herd approximately 7–14 days before the onset of the first clinical signs on 1 January 2025. It therefore seems most likely that the virus was introduced to the herd in the second half of December 2024.

Between 1 December 2024 and the detection of FMD (10 January 2025), about 200,000 animals were transported to or from farms in Berlin and Brandenburg. More than 75% of them were pigs, followed by cattle, sheep, and goats (see **Supplemental Table S3**). Within the analysed trade network, there were 1,939 farms, including 1,366 cattle farms, 199 pig farms, 121 sheep farms, 22 goat farms and 231 mixed farms (keeping more than one susceptible species). Most of the farms were trading only within their species, resulting in separated networks. The number of transports of cattle was much higher (1,707) than of pigs (264) or sheep and goats (137 and 30, respectively). The network was clustered into small units, and the largest giant strongly connected component (GSCC) included only 17 farms (see **Supplemental Figure S5a**). Overall, the analysis showed that there were few movements over the investigated period, particularly in the trade network of the farms trading to and from the later restricted zones (see **Supplemental Figure S5b**).

A simulation model showed that the duration of the epizootic and the number of infected herds was lower in simulations using depopulation within the 1 km zone in addition to culling only the infected herds themselves. Overall, in 50% of the simulation runs, the number of infected herds was between 1 and 16. When deliberately selecting the affected buffalo herd as the index herd in the simulation, the number of affected herds was even lower and, in many simulation runs, *only* the index herd was affected (23.7% of all runs in the depopulation scenario) (**Figure 6**). The model results support the observation that an introduction of FMD into this particular buffalo herd does not necessarily lead to any additional outbreaks.

**Figure 6:**
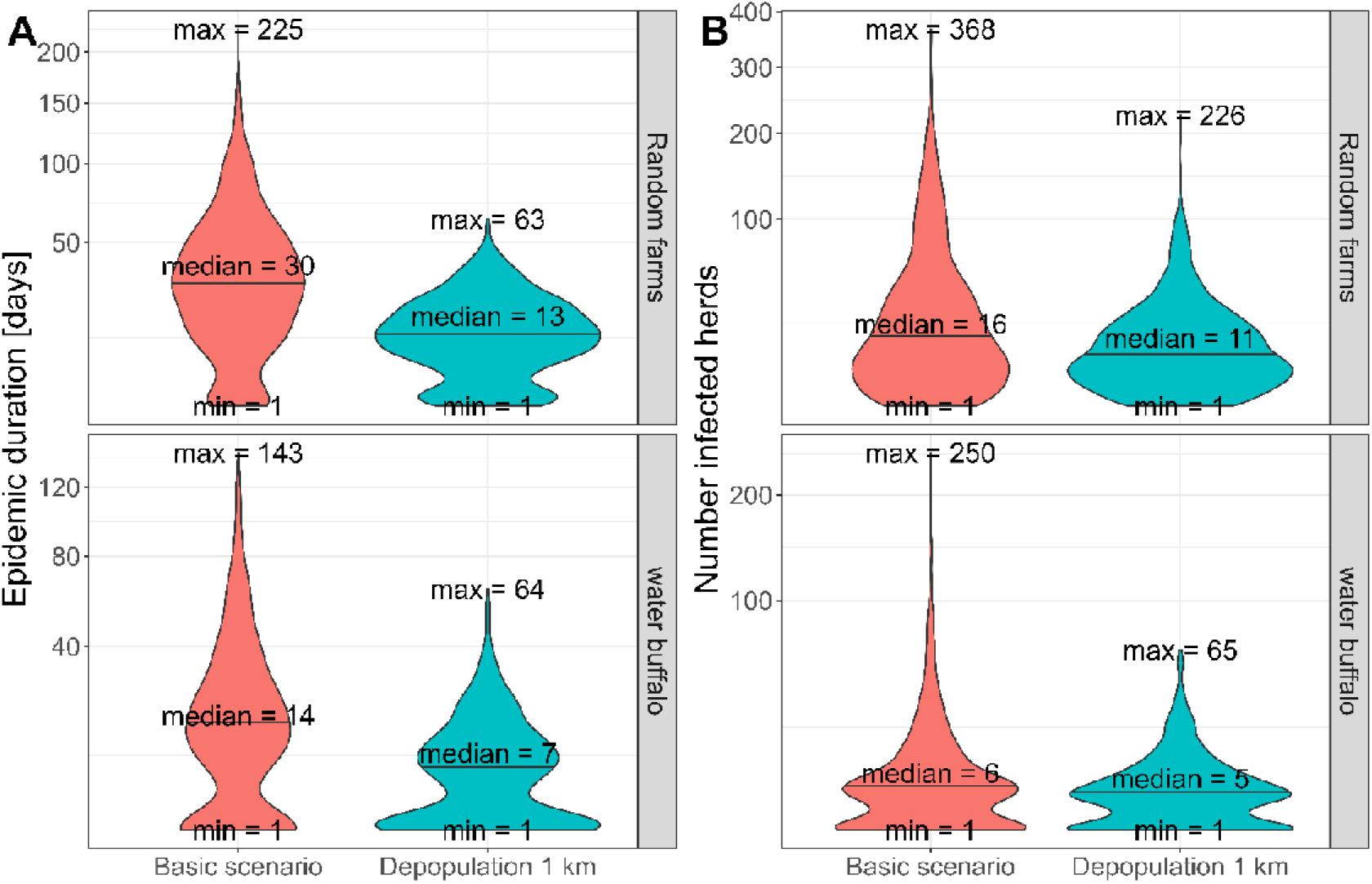
Modelling of the epizootic duration in different scenarios. **a** The epizootic duration when selecting a random herd within Brandenburg or Berlin (upper part) or deliberately selecting the affected water buffalo herd (lower part) as the index herd. **b** the number of infected herds over the course of the simulated epizootic, when selecting a random herd within Brandenburg or Berlin (upper part) or deliberately selecting the affected water buffalo herd (lower part) as the index herd. The epizootic was modelled using 1000 simulation runs and applying either of two control strategies (“basic scenario”: culling of the infected herds only, shown in red; and “depopulation 1 km”: culling of infected herds and surrounding herds with susceptible animals within 1 km, shown in turquoise).

### Introduction hypotheses

FMDV is most easily spread by direct contact between infected and susceptible animals, but contaminated materials (clothing, agricultural equipment, transport vehicles) and feed (including untreated swill) can also be a source of contagion^1^. In the case of the infected farm in Brandenburg, no animals had been introduced into the herd since 2021, there had been no artificial insemination, no equipment or vehicles were shared with any other holding, there had been no visitors in contact with the animals and the persons caring for the animals had not travelled to any country where FMD is known to occur. All feed came from the farm itself or from the immediate vicinity. Introduction by contact with infected wild animals is unlikely since no evidence of FMDV circulation in local wildlife has been found (see below). The owner of the buffalo had previously observed roe deer visiting the pasture and accessing the feed and salt licks, but no wild boar.

Ultimately, two hypotheses remain unrefuted: accidental exposure to contaminated animal products (i.e., food waste) or a deliberate introduction of infected material into the herd. It was reported by the owner that crows had been taking food scraps from nearby waste receptacles and scattering them across the pasture. The outbreak farm is located near a major highway and close to Berlin, making the area a popular destination for walkers and dog owners from the city. Berlin’s strong international ties may facilitate human-associated introductions of FMDV from affected regions. The ad-hoc importation of meat, milk and their products by returning travellers in their personal luggage is restricted by EU legislation^11^, but illegal imports still occur and often remain undetected^12^. Oral exposure to contaminated feed is unlikely to cause infection in ruminants^1^, but it is not impossible. In the absence of any concrete evidence for either hypothesis, however, it appears unlikely that the exact route of introduction will ever be resolved.

### Outbreak resolution

In accordance with EU animal disease control regulations^13^, restricted zones were immediately implemented surrounding the affected farm (**Figure 5**). A standstill for all movements of susceptible animals was maintained for the whole territory of Brandenburg and the whole territory of Berlin until 17 and 27 January 2025, respectively. All holdings with susceptible animals in the 3-km protection zone that had not been depopulated (8 holdings, 94 animals) were visited by official veterinarians three times (7 days apart). All animals were clinically examined, and nasal swabs and serum samples were taken repeatedly. All holdings with susceptible animals in the 10-km surveillance zone (182 holdings, 6,611 animals) and 6 contact holdings outside of the zones (2,887 animals) were visited by official veterinarians twice (14 days apart). The overall number of holdings with susceptible animals in the restricted zones was 193. The animals were clinically examined, and swabs and serum samples were taken repeatedly. In holdings with fewer than 50 animals, all were examined and sampled; in holdings with more than 50 animals, enough animals were examined and sampled to detect a 2% disease prevalence with 95% confidence. Swabs and serum were also collected from susceptible wild animals hunted in the restricted zones, comprising about 500 animals from 1 December 2024 to 31 March 2025. The swabs were tested by FMDV RT-qPCR and the sera were tested with an antibody ELISA. No FMDV RNA or specific antibodies were found in any animal other than the buffalo in the index herd.

Several thousand susceptible animals that had been shipped from Brandenburg to other parts of Germany (n = 5681) and the EU in December and early January were also tested at their destinations, but all were negative.

### Concluding remarks

The European Commission for the Control of Foot-and-Mouth Disease (EuFMD), which also supports the global strategy for progressive control of FMD, gave valuable advice to German authorities concerning preventive and control measures during the event and beyond.

A containment zone as defined in article 4.4.7 in conjunction with article 8.8.10 of the WOAH Terrestrial Animal Health Code^14^ with a roughly 6 km radius around the infected farm (comprising 36 holdings with susceptible animals) was established^15^ and approved by WOAH effective 12 March 2025, with the rest of Germany regaining its FMD-free status upon recognition of the containment zone by WOAH. This containment zone was then lifted in April 2025, three months after the killing and disposal of the buffalo herd. The status of a “FMD-free country where vaccination is not practiced” was reinstated by WOAH for the entirety of Germany on 14 April 2025.

## METHODS

### Histopathology, immunohistochemistry and RNA *in situ* hybridization

Formalin-fixed, paraffin-embedded tissues were processed for routine haematoxylin and eosin (H&E) staining. Immunohistochemistry was performed on consecutive lung and heart sections using the avidin-biotin-peroxidase complex (ABC) method. Briefly, sections were dewaxed in xylene, followed by rehydration in descending graded alcohols. Endogenous peroxidase was quenched with 3% hydrogen peroxide in distilled water for 10 minutes at room temperature (RT). Antigen heat retrieval was performed using citrate buffer (pH 6.0) in a steamer for 20 minutes followed by a cooling period. Nonspecific antibody binding was blocked by normal goat serum (1:2 diluted in TBS) for 30 minutes at RT. The primary antibody (in-house polyclonal rabbit anti-FMDV serum, “MKS-3ABC-3D”) was applied for 1 h at RT (1:1000, diluted in TBS), the secondary biotinylated goat anti-rabbit antibody (Vector Laboratories, USA; 1:200 diluted in TBS) was applied for 30 minutes at RT. Colour was developed by incubating the slides with ABC solution (Vectastain Elite ABC kit; Vector Laboratories), followed by exposure to 3-amino-9-ethylcarbazole substrate (AEC; Dako, USA). The sections were counterstained with Mayer’s haematoxylin and coverslipped. Archived tissue slides from experimental FMD studies served as positive controls. As a negative control, sections were tested with rabbit pre-immune serum instead of MKS-3ABC-3D.

RNA *in situ* hybridization was performed on consecutive lung and heart sections using custom-designed RNAScope probes against the highly conserved FMDV non-structural protein 3D (Advanced Cell Diagnostics, USA). The RNAScope 2-5 HD Reagent Kit-Red (Advanced Cell Diagnostics) was applied according to the manufacturer’s instructions. As technical assay controls, a positive control probe (peptidylprolyl isomerase B) and a negative control probe (dihydrodipicolinate reductase) were included. Archived tissue slides from experimental FMD studies served as positive controls.

All sides were scanned using a Hamamatsu S60 scanner and evaluation was done using the NDPview.2 plus software (Version 2.8.24; Hamamatsu Photonics, Japan) by a board-certified pathologist.

### Nucleic acid extraction and FMDV real-time RT-qPCR

At the state veterinary diagnostic laboratory, RNA was extracted from 200 µl of serum, swab fluid or tissue macerates (lung, spleen, tonsil, lymph node, intestine) with the innuPREP AniPath DNA/RNA Kit 2.0 KFFLX (iST Innuscreen GmbH, Germany). One-step RT-qPCR was performed using primers and a TaqMan probe targeting the 3D coding region^16^ and the internal ribosomal entry site^17^ of FMDV using AgPath-ID One-Step RT-PCR reagents (Thermo Fisher Scientific, Germany). At the NRL, RNA was extracted from 140 µl of serum, swab fluid or the supernatant of freshly prepared tissue macerates with the Viral RNA Mini kit (Qiagen, Germany) and used for FMDV one-step RT-qPCRs as described above.

### Serology

Antibodies against non-structural proteins (NSP) of FMDV were detected in serum samples with the ID Screen FMD NSP Competition ELISA (Innovative Diagnostics, France). After the serotype of the outbreak virus had been identified, the ID Screen Type O Competition ELISA (Innovative Diagnostics) was used to detect antibodies against structural proteins.

### Virus isolation, serotype determination and vaccine matching

Tissue samples of the third dead buffalo were collected during the post-mortem examination. A piece of lung tissue (approximately 10 mm^3^) was macerated in 1 ml of phosphate-buffered saline using a 5-mm steel bead in a TissueLyser II instrument (Qiagen) for 2 minutes at 30 Hz. The macerate was centrifuged at high speed for 1 minute and 100 µl of supernatant were added to a confluent monolayer of recombinant porcine kidney cells expressing bovine αVβ6 integrin^18^ in a 25 cm^2^ culture flask. Strong cytopathic effect was evident after 30 hours of incubation at 37°C. The cell lysate was clarified by centrifugation and the supernatant was stored at -80°C. FMDV of serotype O was detected in the culture supernatant by a standard double-antibody sandwich ELISA^19^ with polyclonal antibodies specific for the 7 serotypes of FMDV.

Vaccine matching for the German field isolate was performed by the European Union reference laboratory for FMD (Anses, Maisons-Alfort) by virus neutralisation tests (VNT) with IB-RS-2 cells as described in the WOAH Manual. The isolate was tested against three bovine vaccinal sera (BVS; provided by Boehringer Ingelheim, Germany) collected 21 days after vaccination. Each BVS was a pool of serum from five cattle vaccinated with either O-3039, O PanAsia-2 or O_1_ Manisa antigen. The r_1_-values shown in **Supplemental Table S2** were calculated by dividing the arithmetic mean of the neutralisation titre of the BVS against the field isolate by the arithmetic mean of the neutralisation titre of the BVS against the homologous vaccine strain. An r_1_-value larger than 0.3 suggests that a high-potency vaccine will be protective against the heterologous field virus. Each r_1_-value is based on three sets of VNT results.

### Genome sequencing

Sample processing for sequencing followed the procedures described by Köster and colleagues^20^. Briefly, RNA was extracted from the aqueous supernatant of homogenized (TissueLyser II, Qiagen) and TRIzol-inactivated organ (lung, intestine, lymph nodes, spleen, tonsil) and blood samples derived from infected water buffalos utilizing the RNAdvance Tissue Kit (Beckman Coulter, Germany) on a KingFisher Flex platform (Thermo Fisher Scientific). After RNA quantification and quality control, up to 350 ng of RNA were used as input for the cDNA synthesis in the Super Script IV First Strand Synthesis System (Thermo Fisher Scientific), followed by a second-strand synthesis step conducted with the NEBNext Ultra II Non-Directional RNA Second Strand Synthesis Module (E6111L, New England Biolabs, Germany). The resulting double-stranded DNA was cleaned and concentrated with a 1.8-fold volume of AMPure XP beads (Beckman Coulter). The cleaned cDNA was subdivided into two fractions to enable parallel library preparation for Oxford Nanopore and Ion Torrent-based sequencing.

For Ion Torrent sequencing, the double-stranded DNA was fragmented with an M220 Focused ultrasonicator (Covaris, USA) and then converted to libraries with the NEBNext Fast DNA Library Prep Set for Ion Torrent (E6270L, New England Biolabs) in combination with IonXpress Barcode Adapters (Thermo Fisher Scientific). Quality control of libraries was performed on a Bioanalyzer 2100 using the High Sensitivity DNA Chip Kit (Agilent Technologies, Germany) and quantities were determined with the QIAseq Library Quant Assay kit (Qiagen). Library pools were prepared and loaded on Ion 530 chips with an Ion Chef instrument (Thermo Fisher Scientific) and sequenced on an S5XL platform (Thermo Fisher Scientific) in 400 bp mode.

For the generation of Nanopore-compatible libraries, 200 ng of cDNA were used in the workflow of the Rapid Barcoding Kit 24 V14 (SQK-RBK114.24, Oxford Nanopore Technologies [ONT]). The obtained library pool was sequenced on an R10.4.1 PromethION Flow Cell (FLO-PRO-114M, ONT) and a P2 Solo sequencing device (ONT) running the high-accuracy model (v4.3.0; 400 bps) for base calling.

### Sequence analysis

The raw Nanopore sequence reads were imported into Geneious and mapped to the FMDV reference genome NC_039210 using Minimap (v2.24; “map-ont” preset)^21^. Only the library prepared from lung tissue yielded sufficient genome coverage and was selected for further analysis. The full genome consensus sequence was deduced and compared to available references using the NCBI BLASTn web service. Reference mapping was then repeated against FMDV reference OP957418.1, the FMDV genome that produced the best BLASTn hit, and the initial consensus sequence was manually inspected and corrected if necessary.

The initial full-length genome sequence was further confirmed and corrected by trimming and mapping the Ion Torrent-derived reads using Newbler software (v3.0, Roche/454 Life Sciences). Finally, the Ion Torrent and Nanopore datasets were *de novo* assembled using Megahit (v1.2.9)^22^ and Flye (v2.9.2-b1786), respectively, and the resulting contigs were matched to the initial consensus sequence using Minimap (v2.24; “map-ont” preset)^21^. The final full FMDV genome sequence of O/GER/BB/1/2025 was annotated and uploaded to GenBank with the accession number PV446528.

### Phylogenetic analysis of VP1

Initially, the VP1-coding region of the German FMDV sequence O/GER/BB/1/2025 was aligned to all 197 available sequences from lineage O/ME-SA/SA-2018 along with representative sequences of the lineages O/ME-SA/Ind-2001 (n=5), O/ME-SA/PanAsia (n=1) and O/ME-SA/PanAsia-2 (n=1) using MUSCLE (v3.8.425)^23^. The sequences were received either from FMDbase at The Pirbright Institute or NCBI Genbank. To assess the overall genetic grouping, a phylogenetic tree was constructed using IQ-TREE (v3.0.1)^24^ running in automatic model selection (-m MFP; ^25^) and applying 10,000 ultrafast (-B 10000)^26^ and SH-aLRT (-alrt 10000) bootstraps each. To assess time-dependent genetic variation in lineage SA-2018, only the 198 VP1 sequences from this lineage were analysed using the same phylogenetic settings as above and the genetic distance (root-to-tip distance) was correlated with the sampling date using TempEst (v1.5.3)^27^. Linear regression and test for significance was done in R (v4.3.1)^28^ using the “lm” function.

Time-aware phylogenetic analyses were performed using BEAST and BEAUti (v1.10.4)^29^ with the VP1 sequence alignment of lineage SA-2018 (n=198). Before the final analysis, we evaluated different combinations of substitution models (“Yang96”, “SRD006”, “GTR+G4+I” and “HKY+G4+I”), clock models (“uncorrelated relaxed clock” and “strict clock”) and tree priors (“GMRF Bayesian skyride” and “constant size”) using the stepping-stone and path sampling methods (path steps: 20; chain length: 200,000). Based on the log marginal likelihood values, we selected the most suitable combination of models and priors and used them for the final analysis (Yang96 + uncorrelated relaxed clock + constant size).

MCMC sampling was performed for 500,000,000 steps, with logs recorded at regular intervals of 10,000 steps. Operators were automatically optimised and outputs included parameter estimates, tree files and population size trajectories. Key statistical operators were checked in the MCMC Trace Analysis Tool (v1.7.2). Specifically, we ensured that the effective sample size of all core model parameters was >400 and that the trace plot had a “fuzzy caterpillar” appearance with no obvious trends. A maximum-clade credibility tree was finally generated using TreeAnnotator (v1.10.4)^29^ and visualized using FigTree (v1.4.5_pre).

### Recombination Analysis

The full genome of the German FMDV sequence O/GER/BB/1/2025 was divided into 500-nt segments, with the trailing 214 nt omitted. Each 500-nt window was used as a query in a MegaBLAST (v2.16.0+)^30^ search against the “core_nt” database (20 June 2025), considering only the top 5 hits from full-length FMDV reference sequences. These full-genome sequences were then aligned with the German FMDV sequence and all available full-genome sequences from lineage SA-2018 using MAFFT (v7.490)^31^. The resulting alignment was analysed for recombination using PhiTest (window size = 100 nt, k = 28) and NeighborNet (GTR distance with empirical frequencies), as implemented in SplitsTree (v4.18.4). The sequences in the alignment were grouped by lineage, after which a BootScan (TN93 distance, window size = 250 nt, step size = 40 nt, bootstraps = 100) was conducted in SimPlot++ (v1.3)^32^, using the German FMDV sequence as the reference sequence. Based on the results, the alignment was split into three parts: (i) 1–4,200 nt, (ii) 4,201–6,125 nt and (iii) 6,126–8,385 nt, which correspond to positions (i) 1–4,046 nt, (ii) 4,048–5,971 nt and (iii) 5,972–8,214 nt in O/GER/BB/1/2025. Individual phylogenetic trees were constructed for each region using IQ-TREE (v3.0.1)^24^, with automatic model selection (-m MFP)^25^ and 1,000 ultrafast bootstraps (-B 1000)^26^.

### Trade network analysis

For the trade network analysis, trade data were used from the German animal traceability and information system (*Herkunftssicherungs-und Informationssystem für Tiere, HI-Tier*). Within the trade network, there were a total of 1,939 farms, including 1,366 cattle farms, 199 pig farms, 121 sheep farms, 22 goat farms, and 231 mixed farms (keeping more than one susceptible species). All movements of cattle, sheep, pig and goats between 1 December 2024 and the detection of FMD on 9 January 2025 were included in the analysis. Individual animal data were used for cattle, but for pigs, sheep and goats, only data on herd level were available.

For each animal or herd, the transports between farms were analysed. The analysis was carried out in R^28^, using the packages “base”, “visNetwork”, “igraph” and “tidyverse”.

### Modelling of disease spread

For modelling the disease spread, an FMD model, which was originally developed by the University of Davis^33^ and modified for Europe, specifically Denmark^34,35^, was used. Data for the model were provided by the affected German federal states (Brandenburg and Berlin) and downloaded from the German animal traceability and information system HI-Tier. A total of 12,107 farms keeping cloven-hoofed animals in Brandenburg and Berlin were included in the model, 89 of which kept buffalo. Data was prepared in R^28^ to fit in the model. Two scenarios were defined: (i) random selection of the index herd among all cloven-hoofed animal herds in Berlin and Brandenburg and (ii) deliberate selection of the affected water buffalo herd in Brandenburg as the index herd. For both scenarios, a standard control strategy (culling only infected herds, no preventive culling) and an extended control strategy with preventive culling of all susceptible animals within 1 km around infected herds were simulated. The model was run for 1,000 times and the results are presented in violin plots using the R package “ggplot2”.

## Acknowledgements

The authors gratefully acknowledge the manifold contributions of Anja Landmesser-Zitzow, Constantin Lorenz, Kira Wisnewski, Karin Pinger, Ulrike Kleinert and Anja Schulz to the diagnostic investigations at the FMD NRL. Silvia Schuparis expertly processed the histopathological samples of buffalo 334.

## Author contributions

Data curation: CS1, DH, FP, JG, KS, LR, NK2, SC. Formal analysis: AB, CS1, FP, JG, ME, NK2, SC. Funding acquisition: CSL, DPK, LBK, MB, SN. Investigation: AB, AR, AR2, CÇ, CS1, CS2, FP, GG, KA, KS, ME, NR, PZ, RM, SC, SK, SW, ÜP, WIB. Methodology: AB, CS1, DH, DPK, FP, JG, ME, NK. Resources: BH, CS1, DH, JG, ME, NK2, SN. Software: CS1, DH, FP, JG, NK2. Supervision: BH, CK, CSL, DH, DPK, LBK, MB, ME, SB, SL1, SN, WIB. Visualization: AB, CS1, FP, JG, ME. Writing – original draft: ME, CS1, FP, KS, CSL, MB. Writing – review & editing: ME, CS1, FP, JG, KS, LR, SB, WIB, CS2, RM, NR, SN, NK, SF, AR, KA, SL1, CH, SL2, BH, SC, DH, PZ, AB, CÇ, ÜP, SK, LMSA, NK2, AR2, GG, LBK, DPK, CK, CSL, MB.

