## Supplemental Material for "Buffaloed in Brandenburg: Germany’s first Brush with Foot-and-Mouth Disease after four Decades of Freedom"

Eschbaumer *et al.* 2025

### Supplemental Figures

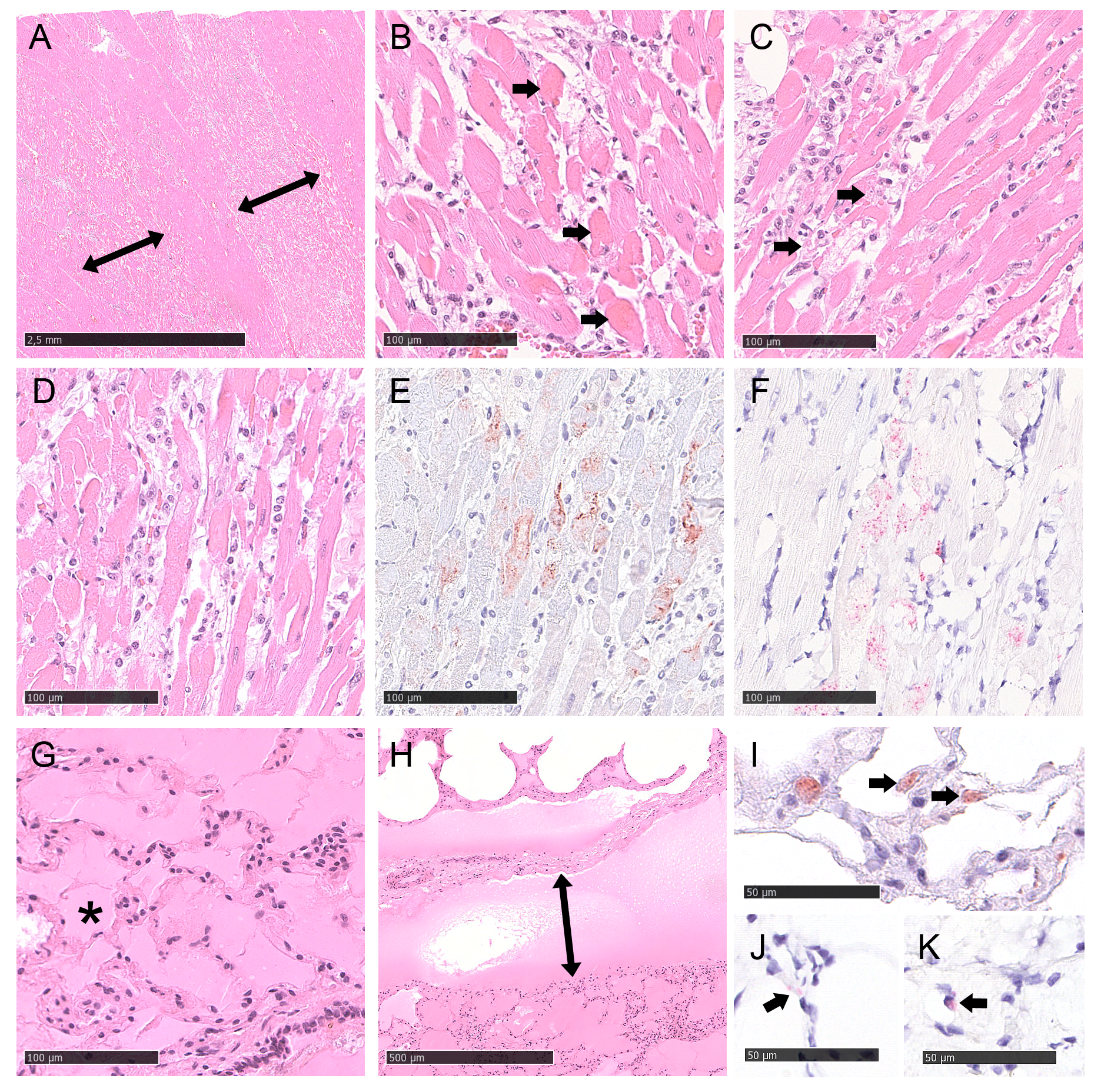

**Supplemental Figure S1: Histological findings.** Histopathology of heart (A-F) and lung (G-K) of buffalo 334 with detection of FMDV antigen (E, I) and RNA (F, J-K). (A) Histologically, similar to the macroscopic picture, there is streaky brightening (↕) of the myocardium. Haematoxylin and eosin (H&E) staining, bar = 2.5 mm. (B) Cardiomyocytes show either hyaline degeneration (→) with glassy, homogeneous cytoplasm, or (C) acute necrosis (→) and appear as cellular debris. Immune cell infiltrates comprise mainly lymphocytes and macrophages and are located between affected cells H&E, bar = 100 µm. (D-E) Cardiomyocytes affected and shown with H&E in (D) exhibit intralesional FMDV antigen as shown by immunohistochemistry (IHC) in (E) using 3-amino-9-ethylcarbazole (AEC) for red-brown chromogen labelling and haematoxylin counterstain and FMDV genome in (F), as shown by RNA in situ hybridization (RNA-ISH) using fast red for labelling and a haematoxylin counterstain, bar = 100 µm. (G) Alveolar and (H) interstitial oedema with presence of eosinophilic, proteinaceous material within the alveoli (*) and expanding the interstitial (↕) space, H&E, bar (G) = 100 µm and (H) = 500 µm. (I) FMDV antigen detection in the cytoplasm of the pulmonary epithelium, IHC, AEC chromogen labelling, haematoxylin counterstain, bar = 50 µm. (J-K) FMDV genome detection in the cytoplasm of very few pulmonary epithelial cells, morphologically consistent with (J) type I and (K) type II pneumocytes. RNA-ISH, fast red labelling, haematoxylin counterstain, bar = 50 µm.

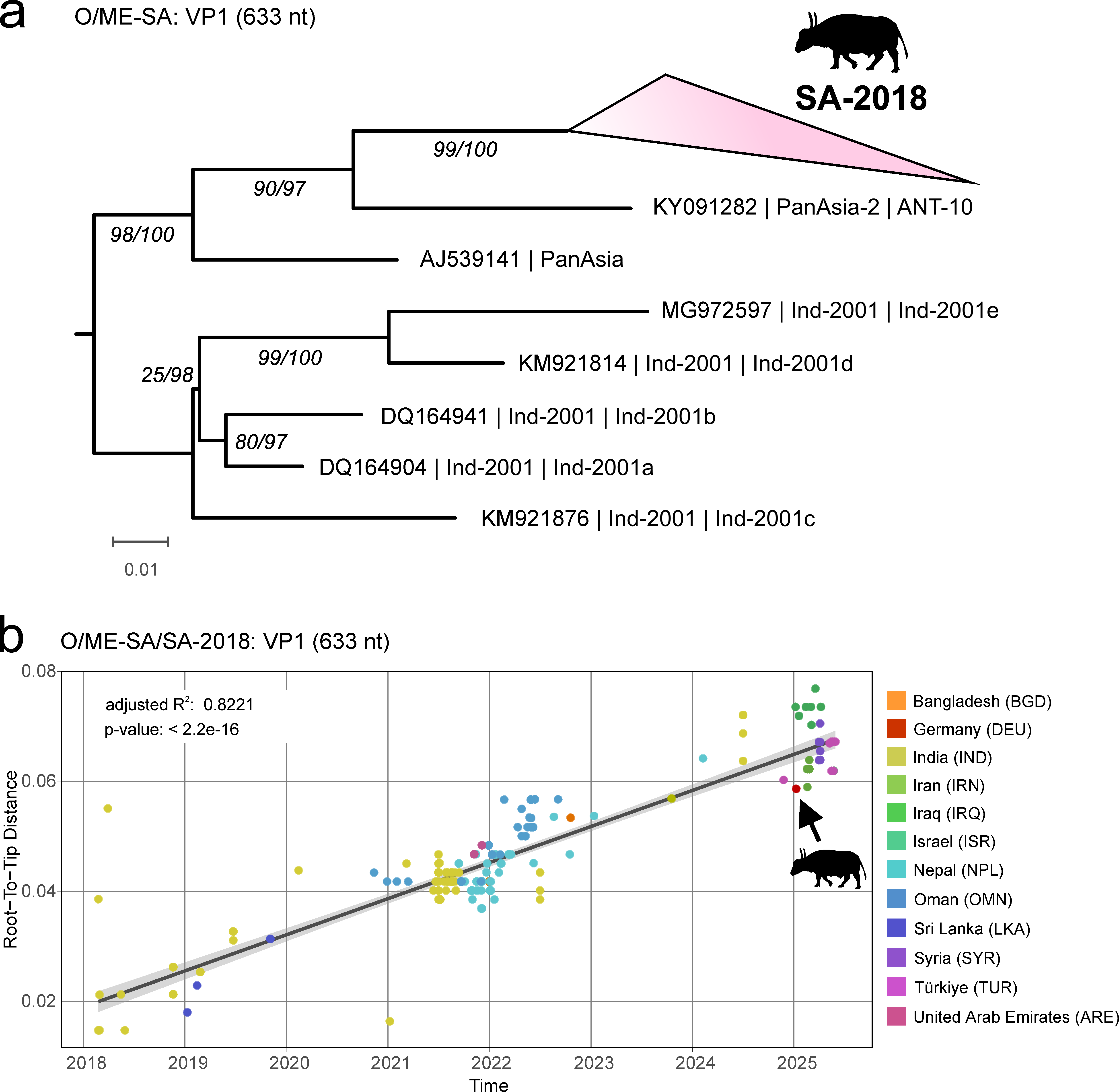

**Supplemental Figure S2: Phylogenetic and temporal analysis of the FMDV VP1-coding region.** **a** A maximum-likelihood phylogenetic tree showing the German FMDV VP1 sequence alongside 197 reference sequences from lineage SA-2018 and representative sequences from lineages Ind-2001, PanAsia and PanAsia-2, all within topotype O/ME-SA. The tree was inferred using IQ-TREE (v3.0.1) with automatic model selection and branch support estimated from each 10,000 ultrafast bootstrap and SH-aLRT replicates. Branch support is shown in italics using the scheme: SH-aLRT support (%) / ultrafast bootstrap support (%). **b** The correlation of genetic divergence (root-to-tip distance) with the sampling date for sequences from lineage SA-2018 (n = 198) was estimated using TempEst (v1.5.3). The linear regression line and coefficient of determination (R²) were calculated in R (v4.3.1) using the 'lm' function.

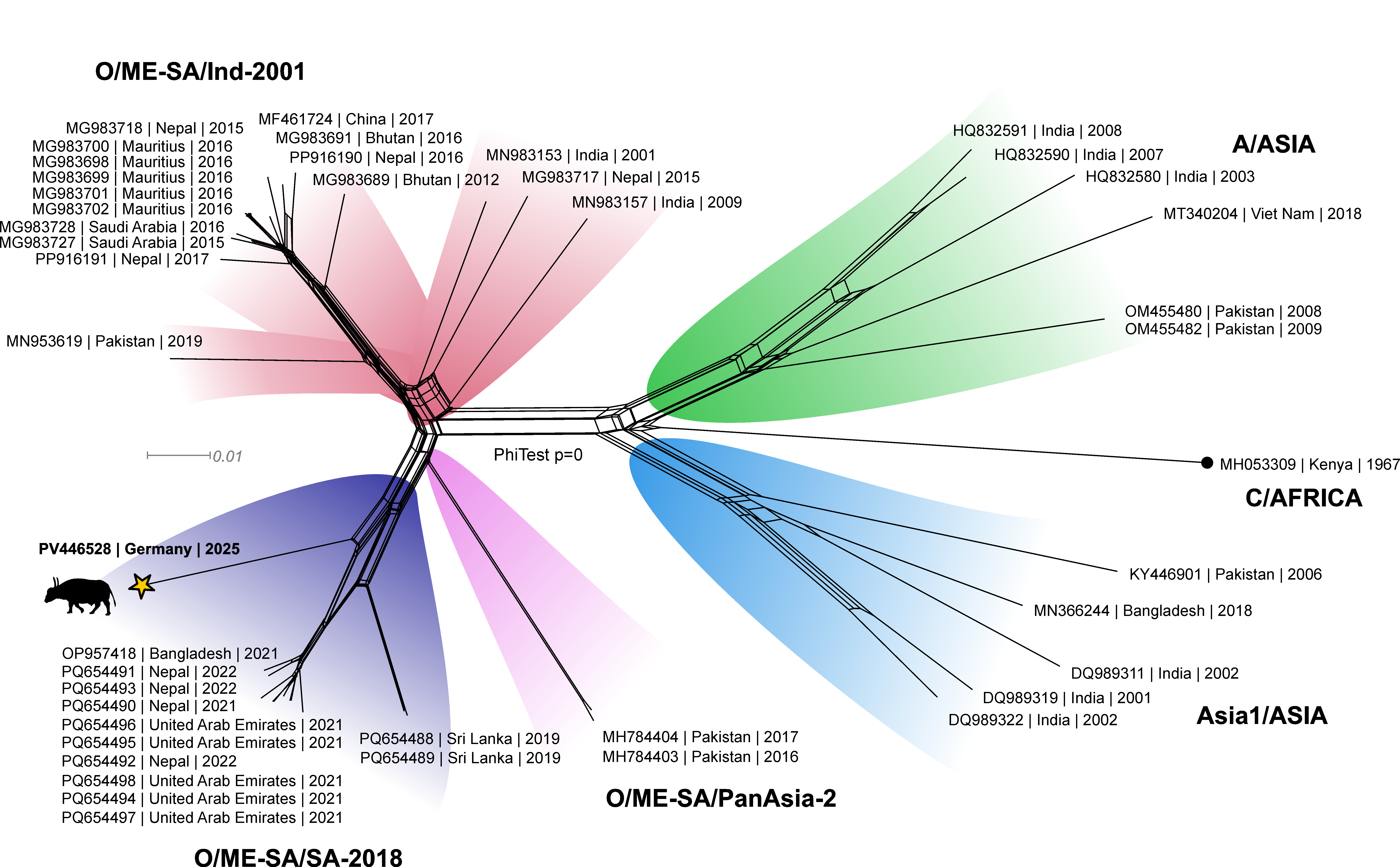

**Supplemental Figure S3: Phylogenetic network of full FMDV genome sequences shows evidence of recombination.**

NeighborNet analysis using an alignment of selected full FMDV genome sequences from different lineages. Phylogenetic conflicting signals, e.g., caused by recombination events, are visualized as box-like structures in the net. The German FMDV strain (star symbol) groups with members of the SA-2018 lineage but its offset position in relation to the other strains indicates recombination. A PhiTest showed statistically significant support for recombination (p-value=0).

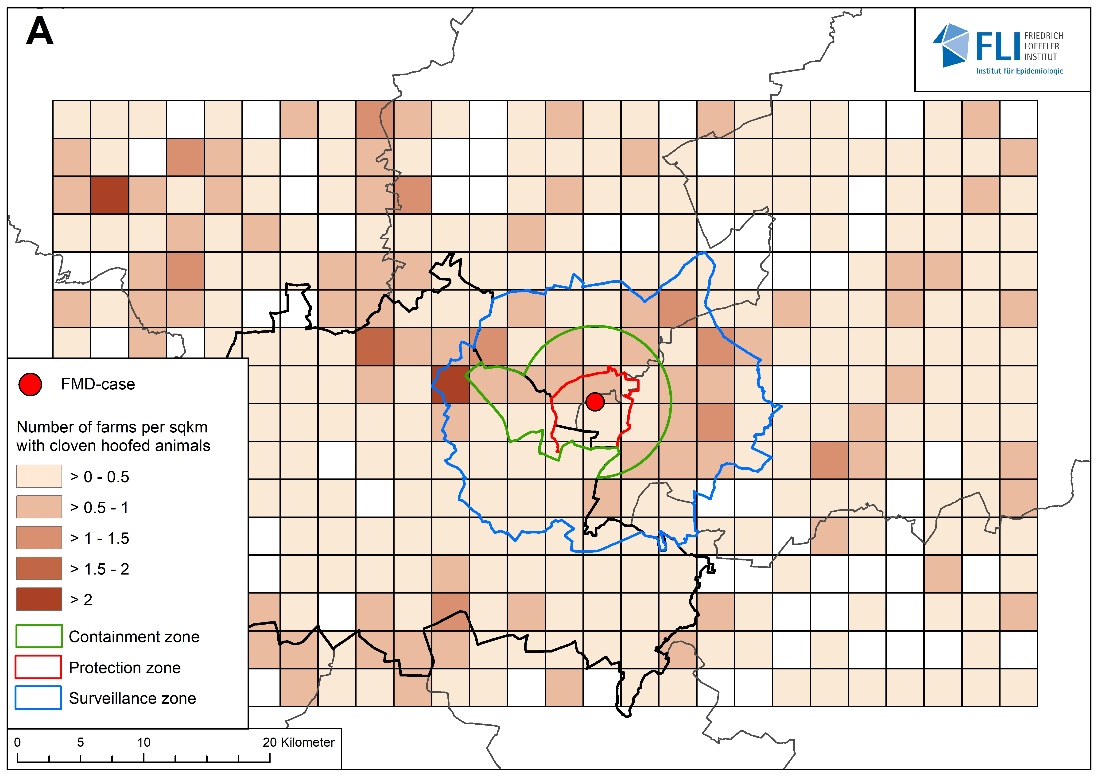

­­

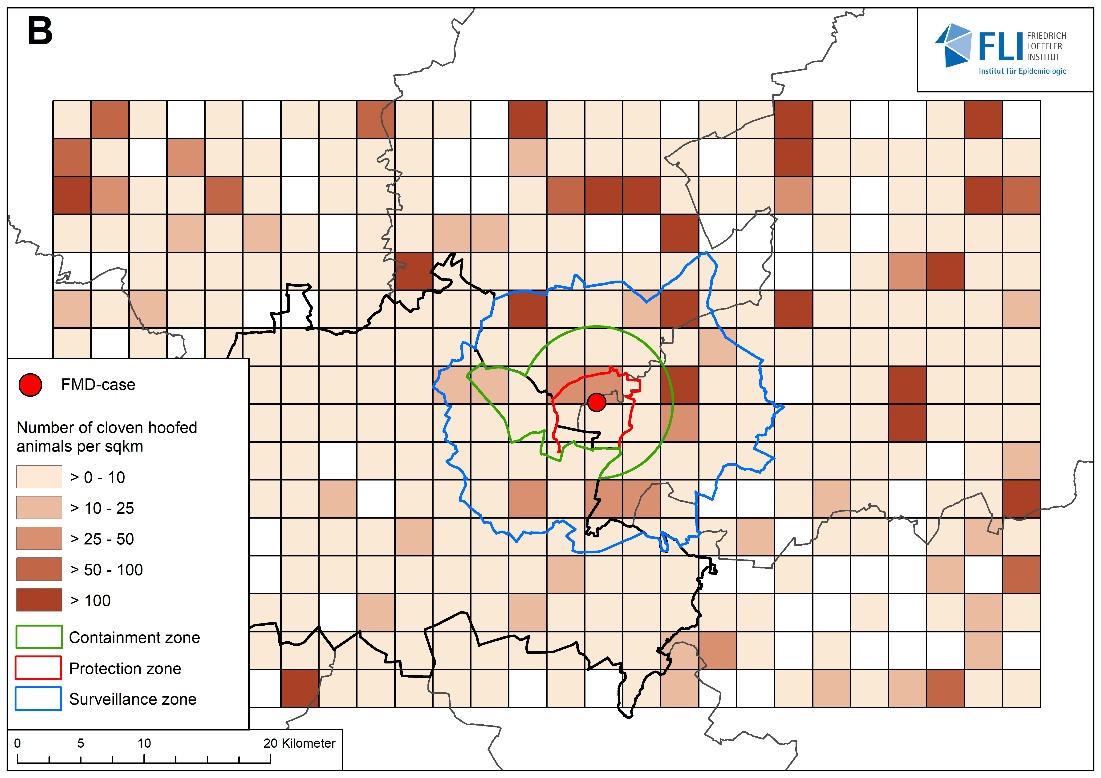

**Supplemental Figure S4: Farm and livestock density.** Density of farms with cloven-hoofed animals (A) and density of cloven-hoofed animals (B) around the outbreak farm. For comparison, the median density of cloven-hoofed livestock in Germany is 42 per km^2^ (Q1–Q3: 19–85).

The red and blue lines show the protection and surveillance zone declared around the outbreak farm on 10 January 2025. The green line shows the containment zone, which was put in place on 12 March 2025.

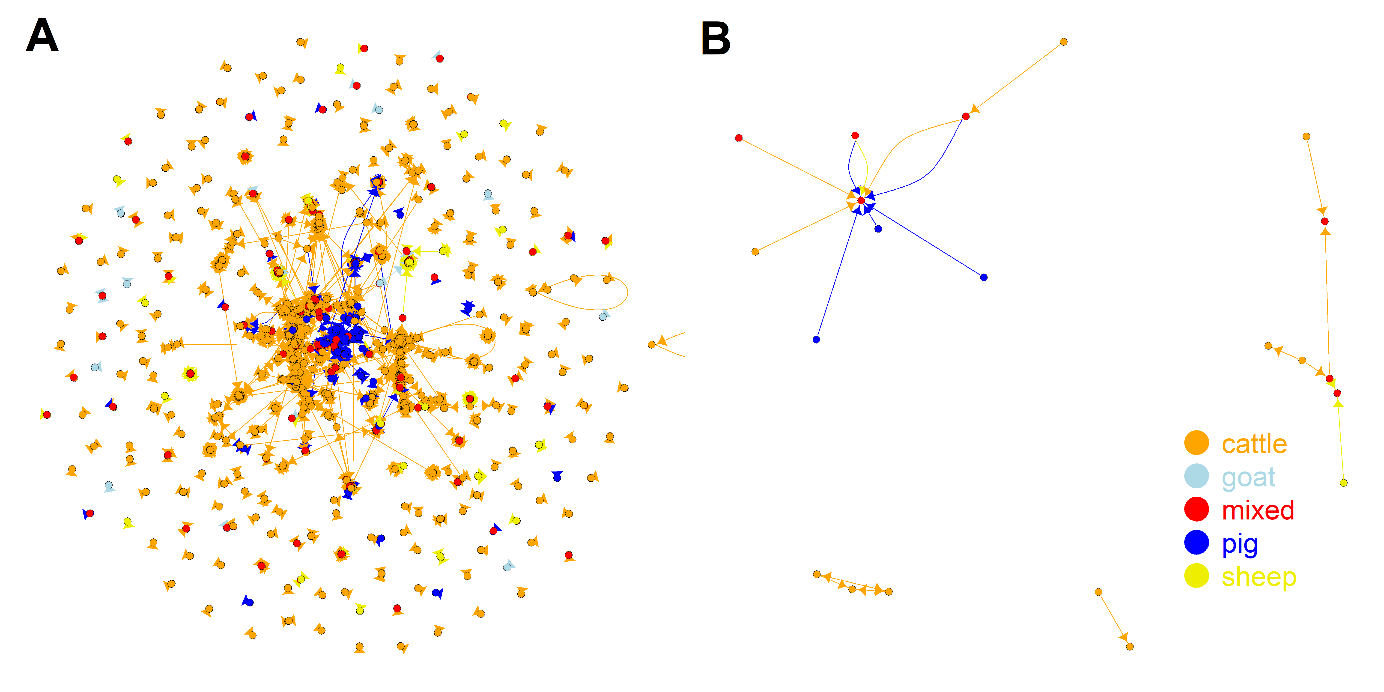
**Supplemental Figure S5: Trade network of animals.** Cattle, swine, goat and sheep movements in (A) Berlin and Brandenburg and (B) the restricted zones from 1 December 2024 to 9 January 2025. Node and edge colours indicate the species of the animals that were transported.

### Supplemental Tables

**Supplemental Table S1: Virological and serological results for 11 culled water buffalo**

Results for the animal (ear tag 334) that had died on the farm on 5 January 2025 and was submitted for necropsy at the state veterinary diagnostic laboratory are not included in this table. No swab samples or lesion material were collected from this animal. The cut-off for the antibody ELISAs is at S/N% 50.

| **animal**  *(ear tag)* | **serum**  *(FMDV Cq)* | **nasal swab**  *(FMDV Cq)* | **mouth lesion**  *(FMDV Cq)* | **foot lesion**  *(FMDV Cq)* | **oldest lesion**  *(age in days)* | **NSP antibody ELISA**  *(S/N%)* | **Type O SP antibody ELISA**  *(S/N%)* |
| --- | --- | --- | --- | --- | --- | --- | --- |
| 168 | 33.5 | 32.8 | none found | 22.6 | not determined | pos (13) | pos (22) |
| 739 | 39.7 | 32.4 | 32.7 | 20.9 | 5 | pos (18) | pos (35) |
| 165 | neg (No Cq) | 37.1 | none found | none found | not determined | pos (31) | pos (22) |
| 565 | neg (No Cq) | 35.8 | 34.0 | none found | >7 | pos (24) | pos (19) |
| 779 | neg (No Cq) | 33.8 | 35.5 | none found | 4-5 | pos (13) | pos (19) |
| 170 | neg (No Cq) | neg | 39.0 | none found | 5 | pos (29) | pos (13) |
| 465 | 35.7 | 37.7 | 36.1 | none found | 5 | pos (27) | pos (22) |
| 740 | neg (No Cq) | 32.4 | 37.8 | none found | not determined | pos (16) | pos (29) |
| 169 | 37.8 | 31.3 | neg (No Cq) | none found | 3 | pos (35) | pos (28) |
| 113 | neg (No Cq) | 33.9 | neg (No Cq) | none found | not determined | pos (19) | pos (26) |
| 167 | neg (No Cq) | 35.2 | neg (No Cq) | none found | not determined | pos (40) | pos (37) |

**Supplemental Table S2: Vaccine matching results**

The r_1_ values shown below represent the one-way serological matches between the vaccine strain and the field isolate, calculated from the comparative reactivity of a pool of bovine sera raised against the vaccine in question, to the vaccine virus and the field isolate. Heterologous neutralisation titres for the field isolate are provided. The result represents the mean of three independent repetitions.

| **Vaccine strain** | **Heterologous titre** | **r_1_ value** |
| --- | --- | --- |
| O-3039 BI | 2.00 | 0.28 |
| O-PanAsia 2 BI | 2.25 | 0.35 |
| O_1_ Manisa BI | 2.15 | 0.63 |

**Supplemental Table S3: Trading activity**

Farm types and species traded to or from farms in Berlin and Brandenburg between 1 December 2024 and 9 January 2025, as reported to the German animal traceability and information system HI-Tier.

| **Farm type** | **Species traded** | **Transports** | **Animals** |
| --- | --- | --- | --- |
| cattle | cattle | 1,562 | 32,211 |
| mixed | cattle | 145 | 2,466 |
| goat | goat | 13 | 77 |
| mixed | goat | 17 | 277 |
| mixed | pig | 57 | 15,550 |
| pig | pig | 207 | 142,307 |
| mixed | sheep | 34 | 3,063 |
| sheep | sheep | 103 | 5,880 |
| **Total:** |  | 2,138 | 201,831 |
